# Tools for *Cre*-mediated conditional deletion of floxed alleles from developing cerebellar Purkinje cells

**DOI:** 10.1101/2024.03.28.587263

**Authors:** Jennifer N. Jahncke, Kevin M. Wright

## Abstract

The Cre-lox system is an indispensable tool in neuroscience research for targeting gene deletions to specific cellular populations. Here we assess the utility of several transgenic *Cre* lines, along with a viral approach, for targeting cerebellar Purkinje cells. Using a combination of a fluorescent reporter line (*Ai14*) to indicate *Cre*-mediated recombination and a floxed *Dystroglycan* line (*Dag1^flox^*) we show that reporter expression does not always align precisely with loss of protein. The commonly used *Pcp2^Cre^* line exhibits a gradual mosaic pattern of *Cre* recombination in Purkinje cells from P7-P14, while loss of Dag1 protein is not complete until P30. *Ptf1a^Cre^* drives recombination in precursor cells that give rise to GABAergic neurons in the embryonic cerebellum, including Purkinje cells and molecular layer interneurons. However, due to its transient expression in precursors, *Ptf1a^Cre^* results in stochastic loss of Dag1 protein in these neurons. *Nestin^Cre^*, which is often described as a “pan-neuronal” *Cre* line for the central nervous system, does not drive *Cre*-mediated recombination in Purkinje cells. We identify a *Calb1^Cre^* line that drives efficient and complete recombination in embryonic Purkinje cells, resulting in loss of Dag1 protein before the period of synaptogenesis. *AAV8*-mediated delivery of *Cre* at P0 results in gradual transduction of Purkinje cells during the second postnatal week, with loss of Dag1 protein not reaching appreciable levels until P35. These results characterize several tools for targeting conditional deletions in cerebellar Purkinje cells at different developmental stages and illustrate the importance of validating the loss of protein following recombination.

**Significance Statement:** The development of *Cre* lines for targeting gene deletions to defined cellular populations has led to important discoveries in neuroscience. As with any tool, there are inherent limitations that must be carefully considered. Here we describe several *Cre* lines available for targeting of cerebellar Purkinje cells at various developmental stages. We use the combination of a *Cre*-dependent fluorescent reporter line and conditional deletion of the synaptic scaffolding molecule Dystroglycan as an example to highlight the potential disconnect between the presence of a fluorescent reporter and the loss of protein.

## Introduction

Over the past thirty-five years, researchers have used transgenic mouse models to define the molecular pathways involved in nervous system development and function. This began with the generation of the first gene knockout mouse lines in 1989 (Joyner et al., 1989; Koller et al., 1989; Navabpour et al., 2020; Schwartzberg et al., 1989; Zijlstra et al., 1989). While the use of knockout mice has led to many fundamental discoveries, the constitutive nature of gene deletion can have inherent limitations. These can include early developmental phenotypes that prevent analysis of a gene’s function at later stages, as well as phenotypes that arise from cellular populations other than the ones being directly studied. This limitation was circumvented by the development of conditional knockouts that enable control over when and where a specific gene is manipulated (Gu et al., 1994; Luo et al., 2020; Tsien, Chen, et al., 1996; Tsien, Huerta, et al., 1996). Spatiotemporal control of gene deletion using the Cre-lox system requires two components: (1) a “floxed” allele of the target gene that incorporates 34 base pair *loxP* recognition sites flanking a critical genomic region of the target gene, and (2) a driver line that expresses *Cre* recombinase under the control of regulatory elements that confer cellular and/or temporal specificity. When *Cre* is expressed in an animal homozygous for the floxed allele, *Cre* drives recombination of the *loxP* sites, deleting the intervening genomic region. Importantly, this deletion only occurs in cells expressing *Cre*, leaving the floxed allele intact in Cre negative cells. For example, one of the original *Cre*-driver lines used regulatory elements of *CamKII*α to drive recombination in specific excitatory neuron populations beginning in the third postnatal week (Tsien, Chen, et al., 1996). This allowed for the investigation of the role of NMDAR1 (GluN1) in hippocampal plasticity and spatial learning when crossed to a *NMDAR1* floxed line, which is lethal when deleted constitutively (Li et al., 1994; Tsien, Huerta, et al., 1996). Similar conditional genetics approaches using different recombinase/recognition sites have been developed, including Flp-FRT (Dymecki & Tomasiewicz, 1998; Park et al., 2011) and Dre-rox (Anastassiadis et al., 2009). The availability of multiple orthogonal recombination systems allows for intersectional approaches that require the co-expression of multiple recombinases to affect gene expression, giving rise to increased cellular specificity.

An important consideration when using conditional deletion is the fidelity of recombination and efficiency of deletion, which is often assayed by crossing the *Cre* line to a *Cre*-dependent reporter line. These reporter lines typically express a fluorescent (*tdTomato*, *EGFP*) or enzymatic (*LacZ*, *Alkaline Phosphatase*) gene from “safe harbor” loci in the genome that are accessible to Cre-mediated recombination. The reporter is preceded by a “lox-stop-lox” (LSL) cassette that prevents constitutive expression in the absence of *Cre*. However, these cassettes can be highly sensitive to low levels or transient expression of *Cre* and therefore can be an insufficient proxy for the deletion of a target floxed allele (Luo et al., 2020). It is therefore imperative to validate the expected deletion of the target allele directly by assaying mRNA to show loss of the expected transcript in Cre positive cells. However, due to differences in the rate of protein turnover, loss of mRNA may not precisely recapitulate the timing of functional protein loss. Therefore, evaluating the loss of protein should define the gold standard for verifying a conditional deletion.

One particular challenge in using conditional deletion strategies for studying developmental events like synapse formation is identifying lines that express *Cre* early in development in a cell-type specific manner. For example, Parvalbumin (PV) positive interneurons in the hippocampus provide are widely studied and commonly targeted by several *PV^Cre^* lines (Hippenmeyer et al., 2005; Madisen et al., 2010). However, *Cre* expression does not initiate in these cells until P10-P12 (Lecea et al., 1995), which is after they begin the process of synaptogenesis around P6 (Doischer et al., 2008). Therefore, the *PV^Cre^* line can be used to examine the role of genes in synaptic maintenance and/or function, but not initial synapse formation. Several lines express *Cre* early in the development of cells that give rise to the hippocampal PV^+^ interneuron population (*Dlx5/6^Cre^*, *GAD67^Cre^*, *Nkx2.1^Cre^*), but these lines target multiple interneuron populations (Potter et al., 2009; Taniguchi et al., 2011).

In this study, we examine the efficiency of several *Cre* lines in driving deletion of the synaptic cell adhesion protein Dystroglycan (Dag1) from cerebellar Purkinje cells. We show that the widely used *Pcp2^Cre^* line is insufficient for driving loss of Dag1 protein before the period of synaptogenesis. We show that *Ptf1a^Cre^*drives recombination of a *tdTomato* reporter allele in all embryonic GABAergic neurons in the cerebellum, yet results in stochastic loss of Dag1 protein from Purkinje cells. We identify a *Calb1-IRES-Cre-D* (*Calb1^Cre^*) line that drives specific and robust recombination in Purkinje cells by P0, resulting in loss of Dag1 protein prior to synaptogenesis. Finally, we show that AAV mediated delivery of *Cre* results in gradual transduction of Purkinje cells with a lag between reporter expression and loss of Dag1 protein on the order of weeks.

## Materials and Methods

### Animal Husbandry

All animals were housed and cared for by the Department of Comparative Medicine (DCM) at Oregon Health and Science University (OHSU), an AAALAC-accredited institution. Animal procedures were approved by OHSU Institutional Animal Care and Use Committee (Protocol # IS00000539) and adhered to the NIH *Guide for the care and use of laboratory animals*. Animal facilities are regulated for temperature and humidity and maintained on a 12-hour light-dark cycle (lights on 6AM-6PM); animals are provided with 24-hour veterinary care and food and water *ad libitum*. Animals were group housed whenever possible and provided with environmental enrichment in the form of extra crinkle paper and a red plastic shelter.

Experiments were performed between Zeitgeber Time (ZT) 0 and ZT12. Animals older than postnatal day 6 (P6) were euthanized by administration of CO_2_ and animals <P6 were euthanized by rapid decapitation. Mice of both sexes were used for all experiments.

### Mouse Strains and Genotyping

The day of birth was designated postnatal day 0 (P0). Ages of mice used for each analysis are indicated in the figure and figure legends. Mice were maintained on a C57BL/6 background. *Dag1^+/-^* mice were generated by crossing a male *Dag1^flox/flox^* mouse to a female *Sox2^Cre^* mouse to generate germline *Dag1*^Δ^*^/+^* mice, hereafter referred to as *Dag1^+/-^* mice; *Dag1^+/-^* offspring lacking the *Sox2^Cre^* allele were thereafter maintained as heterozygotes. For all *Dag1* conditional crosses, a *Cre* positive *Dag1^+/-^*breeder was crossed to a *Dag1^flox/flox;tdT/tdT^* breeder to generate *Cre* positive *Dag1^flox/+;tdT/+^* controls and *Cre* positive *Dag1^flox/-;tdT/+^* conditional knockouts. Genomic DNA extracted from toe or tail samples using the HotSHOT method (Truett et al., 2000) was used to genotype animals. Primers for genotyping can be found on the Jackson Labs webpage or originating article (see **Table 1**). *Dag1^+/-^*mice were genotyped with the following primers: CGAACACTGAGTTCATCC (forward) and CAACTGCTGCATCTCTAC (reverse). For each mouse strain, littermate controls were used for comparison with mutant mice.

**Table 1.**
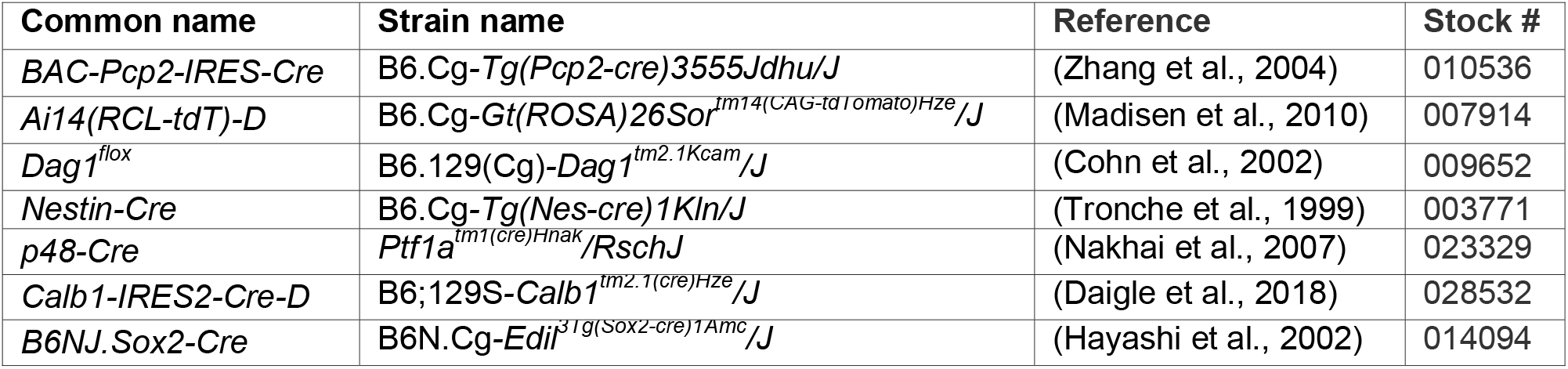
Mouse strains.

### Intracerebroventricular virus injection

Borosilicate capillary glass (World Precision Instruments, Cat. No. 1B150F-4) was pulled into a fine tip and then beveled to an angled point. The capillary was then filled with AAV8-CMV.III.eGFP-Cre.WPRE.SV40 with a titer of 1.00E+13 (Penn Vector Core, University of Pennsylvania) diluted 1:10 in PBS with Fast Green FCF (Fisher Scientific, Cat. No. BP123-10) for visualization (1.00E+12 final titer after dilution). P0 mice were deeply anesthetized through indirect exposure to ice until unresponsive to light touch. An ethanol wipe was used to sterilize the skin of each pup prior to injection. The capillary tip was manually guided into the lateral cerebral ventricle and a Toohey Spritzer Pressure System IIe (Toohey Company, Fairfield, NJ, USA) delivered a 30 psi pulse for 30 msec. The process was repeated in the other hemisphere. Pups then recovered on a warm pad until mobile and returned to their home cage.

### Perfusions and tissue preparation

Brains from mice younger than P21 were dissected and drop fixed in 5 mL of 4% paraformaldehyde (PFA) in phosphate buffered saline (PBS) overnight for 18-24 hours at 4°C. Mice P21 and older were deeply anesthetized using CO_2_ and transcardially perfused with ice cold 0.1M PBS followed by 15 mL of ice cold 4% PFA in PBS. After perfusion, brains were post-fixed in 4% PFA for 30 minutes at room temperature. Brains were rinsed with PBS, embedded in 4% low-melt agarose (Fisher cat. no. 16520100), and sectioned at 70μm using a vibratome (VT1200S, Leica Microsystems Inc., Buffalo Grove, IL) into 24-well plates containing 1 mL of 0.1M PBS with 0.02% Sodium Azide.

### Immunohistochemistry

Free-floating vibratome sections (70μm) were briefly rinsed with PBS, then blocked for 1 hour in PBS containing 0.2% Triton-X (PBST) plus 2% normal goat or donkey serum. For staining of Dystroglycan synaptic puncta, an antigen retrieval step was performed prior to the blocking step: sections were incubated in sodium citrate solution (10mM Sodium Citrate, 0.05% Tween-20, pH 6.0) for 12 min at 95°C in a water bath followed by 12 min at room temperature. Free-floating sections were incubated with primary antibodies (**Table 2**) diluted in blocking solution at 4°C for 48-72 hours. Sections were then washed with PBS three times for 20 min each. Sections were then incubated with a cocktail of secondary antibodies (1:500, Alexa Fluor 488, 546, 647) in blocking solution containing Hoechst 33342 (1:10,000, Life Technologies, Cat. No. H3570) overnight at room temperature followed by three final washes with PBS. Finally, sections were mounted on Fisher Superfrost Plus microscope slides (Fisher, Cat. No. 12-550-15) using Fluoromount-G (SouthernBiotech, Cat. No. 0100-01), covered with #1 coverslip glass (Fisher, Cat. No. 12-541-025), and sealed using nail polish.

**Table 2.**
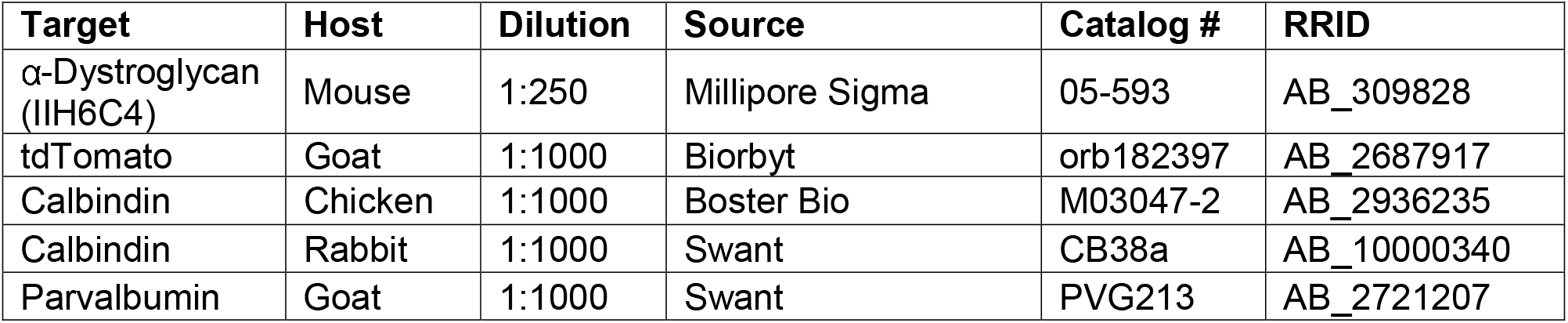
Primary antibodies used for immunohistochemistry.

| Target | Host | Dilution | Source | Catalog # | RRID |
| --- | --- | --- | --- | --- | --- |
| α-Dystroglycan (IIH6C4) | Mouse | 1:250 | Millipore Sigma | 05-593 | AB_309828 |
| tdTomato | Goat | 1:1000 | Biorbyt | orb182397 | AB_2687917 |
| Calbindin | Chicken | 1:1000 | Boster Bio | M03047-2 | AB_2936235 |
| Calbindin | Rabbit | 1:1000 | Swant | CB38a | AB_10000340 |
| Parvalbumin | Goat | 1:1000 | Swant | PVG213 | AB_2721207 |

### Microscopy

Imaging was performed on either a Zeiss Axio Imager M2 fluorescence upright microscope equipped with an Apotome.2 module or a Zeiss LSM 980 laser scanning confocal build around a motorized Zeiss Axio Observer Z1 inverted microscope with a Piezo stage. The Axio Imager M2 uses a metal halide light source (HXP 200 C), Axiocam 506 mono camera, and 10X/0.3 NA EC Plan-Neofluar, 20X/0.8 NA Plan-Apochromat objectives. The LSM 980 confocal light path has two multi-alkali PMTs and two GaAsP PMTs for four track imaging. Confocal images were acquired using a 63X/1.4 NA Plan-Apochromat Oil DIC M27 objective. For some experiments utilizing the LSM 980 confocal, a linear Wiener filter deconvolution step (Zeiss LSM Plus) was used at the end of image acquisition with 1.2X Nyquist sampling (**Figure 3 B-C**; **Figure 7 C-E**). Z-stack images were acquired and analyzed offline in ImageJ/FIJI (Schindelin et al., 2012). Images within each experiment were acquired using the same microscope settings. Brightness and contrast were adjusted identically across samples in FIJI to improve visibility of images for publication. Figures were composed in Adobe Illustrator 2023 (Adobe Systems), with graphs assembled in R (Version 4.2.3).

## Results

### *Pcp2^Cre^* drives gradual *Cre*-mediated recombination in Purkinje cells

The simple, well-defined, and stereotyped circuitry of the cerebellum makes it an ideal model system for studying synapse development and maintenance (**Figure 1**). At the center of the circuit are Purkinje cells (PCs), which are the primary output neurons of the cerebellum and project their axons to the deep cerebellar nuclei (DCN) and the vestibular nuclei in the brainstem. Purkinje cells receive excitatory inputs from two different sources. Parallel fibers originate from cerebellar granule cells, the most numerous neuron type in the brain, and provide a large number of weak excitatory inputs to the dendrites of Purkinje cells (Itō, 1984; Palay & Chan-Palay, 2012). Climbing fibers originate from excitatory neurons in the inferior olive, and their axons wrap around the primary dendritic branches of Purkinje cells, forming strong excitatory contacts (Itō, 1984; Palay & Chan-Palay, 2012). In mouse, Purkinje cells initially receive inputs from multiple climbing fibers, which undergo activity-dependent pruning during the first 3 weeks of postnatal development until a 1:1 ratio is achieved (Bosman et al., 2008; Bosman & Konnerth, 2009; Crepel et al., 1976, but see Busch & Hansel, 2023). These inputs represent one of the best-studied examples of synaptic competition in the central nervous system (CNS). Purkinje cells receive the majority of their inhibitory inputs from two types of Molecular Layer Interneurons (MLIs): Basket Cells (BCs) and Stellate Cells (SCs) (Itō, 1984; Palay & Chan-Palay, 2012). BCs form inhibitory contacts on the soma and proximal dendrites of Purkinje cells, whereas SCs innervate the distal dendrites. Each BC/SC contacts multiple Purkinje cells in the same sagittal plane. There are also recurrent inhibitory connections between Purkinje cells (Altman, 1972; Bernard & Axelrad, 1993; Witter et al., 2016).

**Figure 1.**
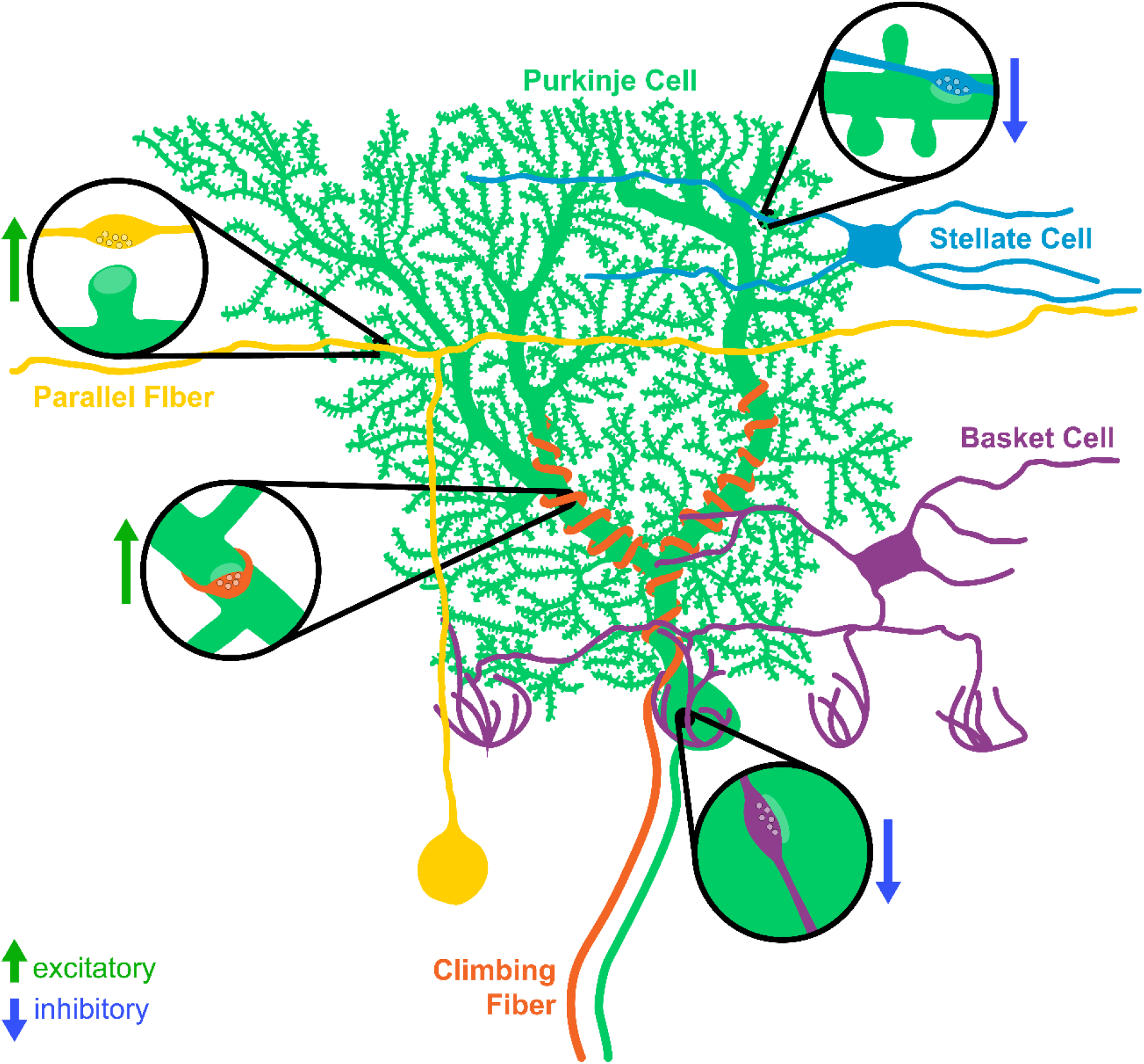
Major synaptic inputs onto Purkinje cells in cerebellar cortex. Purkinje cells (green) receive excitatory inputs from two populations: parallel fibers originating from granule cells (yellow) and climbing fibers originating from the inferior olive (orange). The molecular layer interneurons (basket cells, purple, and stellate cells, blue) provide inhibitory input onto Purkinje cells.

Purkinje cells in the mouse are born between embryonic days 11-13 and elaborate their dendrites over the first four postnatal weeks (Altman, 1972; Miale & Sidman, 1961; Yuasa et al., 1991). They begin to receive excitatory and inhibitory inputs after the first postnatal week and the development of the circuit is largely complete by postnatal day 28 (P28) (Altman, 1972; Kapfhammer, 2004). The most commonly used *Cre* lines for targeting Purkinje cells are the *Pcp2^Cre^* (Zhang et al., 2004) and *L7^Cre^* (Barski et al., 2000) lines. *Pcp2* (*Purkinje cell protein 2,* previously referred to as *L7*), is exclusively expressed by Purkinje cells in the brain, making the *Pcp2* promoter a logical driver for Purkinje cell *Cre* expression. Both the *Pcp2^Cre^* and *L7^Cre^* transgenic lines were generated with similar bacterial artificial chromosome (BAC) constructs and were named to reflect the *Pcp2/L7* gene nomenclature at the time of generation (Barski et al., 2000; Lewis et al., 2004; Zhang et al., 2004). A key limitation of both lines is that *Cre* expression turns on gradually during the period of Purkinje cell development. To rigorously and precisely define the period of *Cre*-mediated recombination in Purkinje cells, we analyzed cerebellar lobule 5/6 from *Pcp2^Cre^;Ai14* brains, which carry a *Cre*-dependent tdTomato reporter knocked in to the *ROSA26* locus (hereafter referred to as *Pcp2^Cre^;ROSA26^LSL-tdTomato^*). The first tdTomato^+^ Purkinje cells were visible at P7 (11.5% of all Purkinje cells), and this increased to 23.9% positive at P9. There was an appreciable increase at P10, with 76.4% of Purkinje cells labeled by tdTomato, and recombination of *tdTomato* was complete by P14 (**Figure 2A, B**). As previously reported, this line shows high specificity for Purkinje cells, as we saw no other tdTomato^+^ neuron types in the cerebellum.

**Figure 2.**
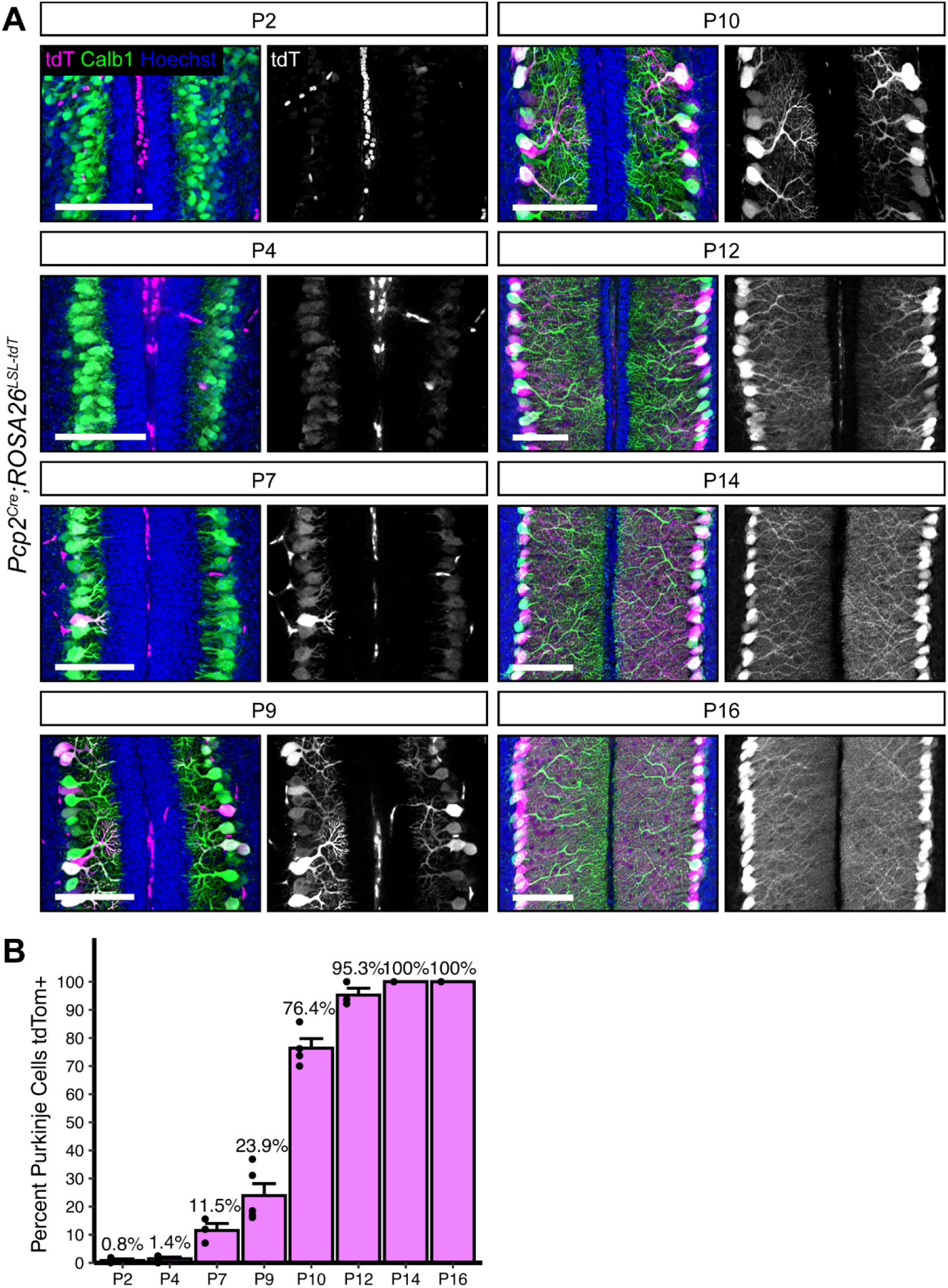
*Pcp2^Cre^*drives *Cre* recombination gradually in the second postnatal week. **(A)** *Pcp2^Cre^* was crossed to a tdTomato reporter to observe the time course of *Cre* recombination. Cerebella were analyzed at time points from P2-P16. *Cre* recombined cells are labeled with tdTomato (magenta). Purkinje cells are visualized with Calbindin immunostaining (green). Nuclei (blue) are most evident in granule cells. Scale bars = 100μm. **(B)** The percent of all Purkinje cells that were tdTomato positive was quantified at each time point.

A key consideration when using *Cre* lines is the timing of protein loss following deletion of floxed alleles. The presence of fluorescent reporters like tdTomato are frequently used as a proxy for *Cre*-mediated recombination, particularly in situations where antibodies for the targeted proteins are lacking. However, this does not account for the timing of mRNA and protein turnover following deletion of the targeted allele, which can vary widely. This is exemplified by Dystroglycan (Dag1), a cell adhesion molecule present postsynaptically at PC:MLI synapses. Dag1 is a stable protein, with a half-life of ∼25 days in skeletal muscle (Novak et al., 2021). Recent work has shown that Dag1 is required for PC:MLI synapse maintenance (Briatore et al., 2020). However, this study was unable to examine its role in synapse formation due to the gradual recombination driven by the *L7^Cre^* line. To resolve Dag1 function in PC:MLI synapse development, we chose to delete *Dag1* with *Pcp2^Cre^*, which recombines with similar timing to *L7^Cre^*, but exhibits increased specificity for Purkinje cells (Barski et al., 2000; Zhang et al., 2004). We generated *Pcp2^Cre^;Dag1^flox/-^* mice (hereafter referred to as *Pcp2^Cre^;Dag1^cKO^*) to see if starting with one allele of *Dag1* already deleted would result in more rapid protein loss. At P16, a timepoint at which all Purkinje cells are tdTomato positive in *Pcp2^Cre^* mice (**Figure 2**), we still observed punctate Dag1 staining consistent with synaptic localization in the majority of *Pcp2^Cre^;Dag1^cKO^* Purkinje cells (**Figure 3A**). Analysis at P30, a period after developmental synaptogenesis is complete, showed a loss of Dag1 protein in *Pcp2^Cre^;Dag1^cKO^* Purkinje cells by immunohistochemistry, which appeared identical at P60 (**Figure 3B**). The incomplete loss of Dag1 protein in *Pcp2^Cre^;Dag1^cKO^*Purkinje cells at P16 highlights a limitation of using a *Cre* line that turns on after gene expression initiates when the protein is particularly stable, as many synaptic molecules are.

**Figure 3.**
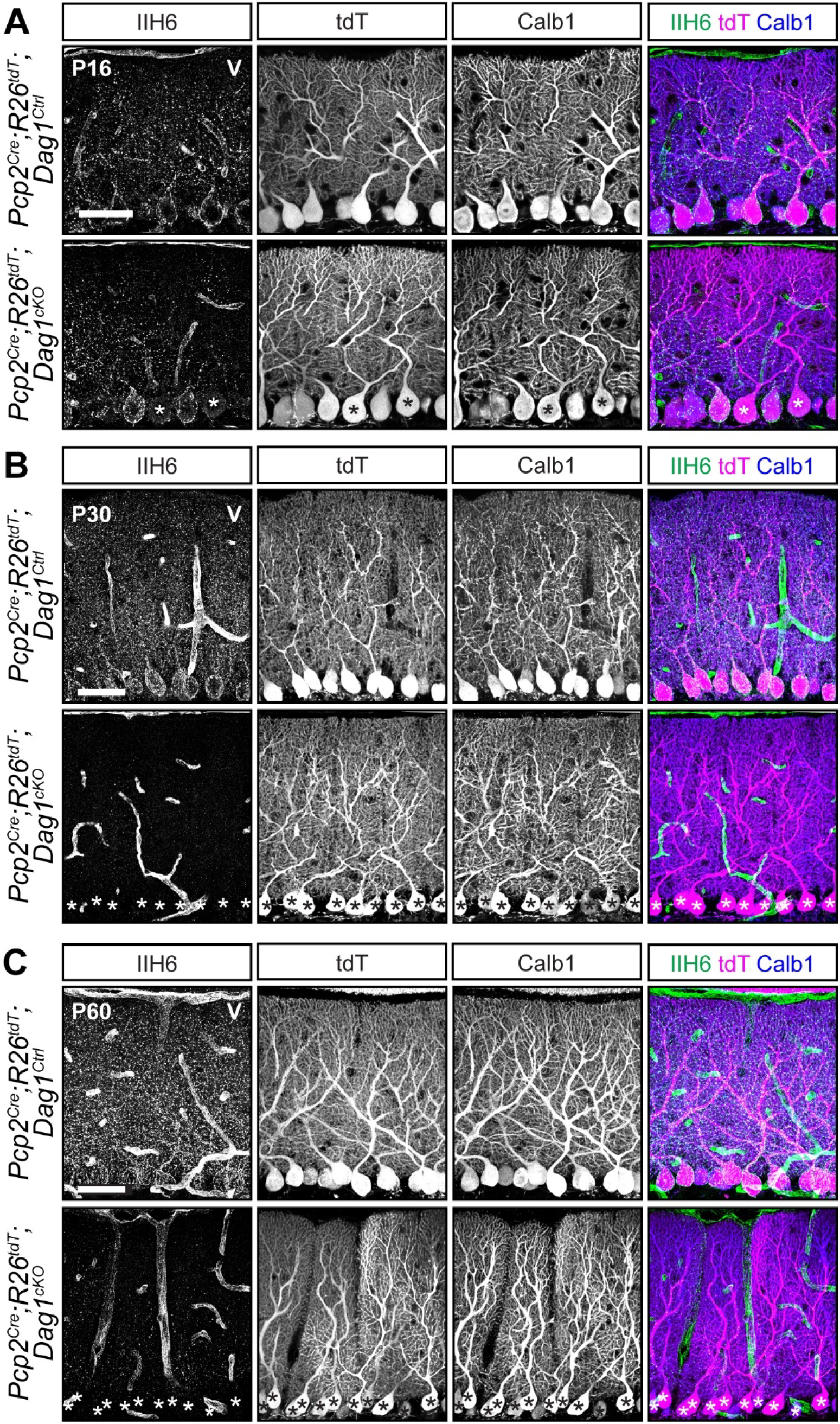
Dystroglycan protein loss lags behind *Pcp2^Cre^* recombination of fluorescent tdTomato reporter. **(A-C)** Lobule V Purkinje cells of *Pcp2^Cre^;ROSA26^LSL-tdTomato^;Dag1^cKOs^*and littermate controls immunostained for α-Dystroglycan (IIH6, green) to visualize Dag1 protein and Calbindin (Calb1, blue) to label Purkinje cells. *Cre* recombination was assessed by native fluorescence of the tdTomato reporter (tdT, magenta). Expression was evaluated at **(A)** P16, **(B)** P30, and **(C)** P60. Asterisks denote Purkinje cells with no detectable IIH6 signal. Scale bars = 50μm.

### Widespread *Nestin^Cre^* recombination does not extend to Purkinje cells

We sought to identify a *Cre* line that would express in Purkinje cells prior to synaptogenesis. We first examined *Nestin^Cre^*, which has widespread expression in the CNS beginning at E10.5 (Graus-Porta et al., 2001; Tronche et al., 1999). Examination of brains from *Nestin^Cre^;ROSA26^LSL-tdTomato^*mice at P8 showed extensive recombination in the forebrain, but minimal recombination in the cerebellum (**Figure 4A**). Higher magnification images confirmed that there was no recombination in Purkinje cells, MLIs, or cerebellar granule neurons at P8 (**Figure 4B**). Consistent with previous reports, there were mild perturbations in cerebellar granule neuron migration in *Nestin^Cre^;Dag1^cKO^* mice due to Dystroglycan’s function in Bergmann glia (**Figure 4C**) (Nguyen et al., 2014). It important to note the particular *Nestin^Cre^*line that we used, B6.Cg-Tg(Nes-cre)1Kln/J (Tronche et al., 1999, RRID: IMSR_JAX:003771), as there are a number of different *Nestin^Cre^* lines that have been generated, some of which have been used to delete genes from Purkinje cells with success (Sun et al., 2014; Takeo et al., 2021).

**Figure 4.**
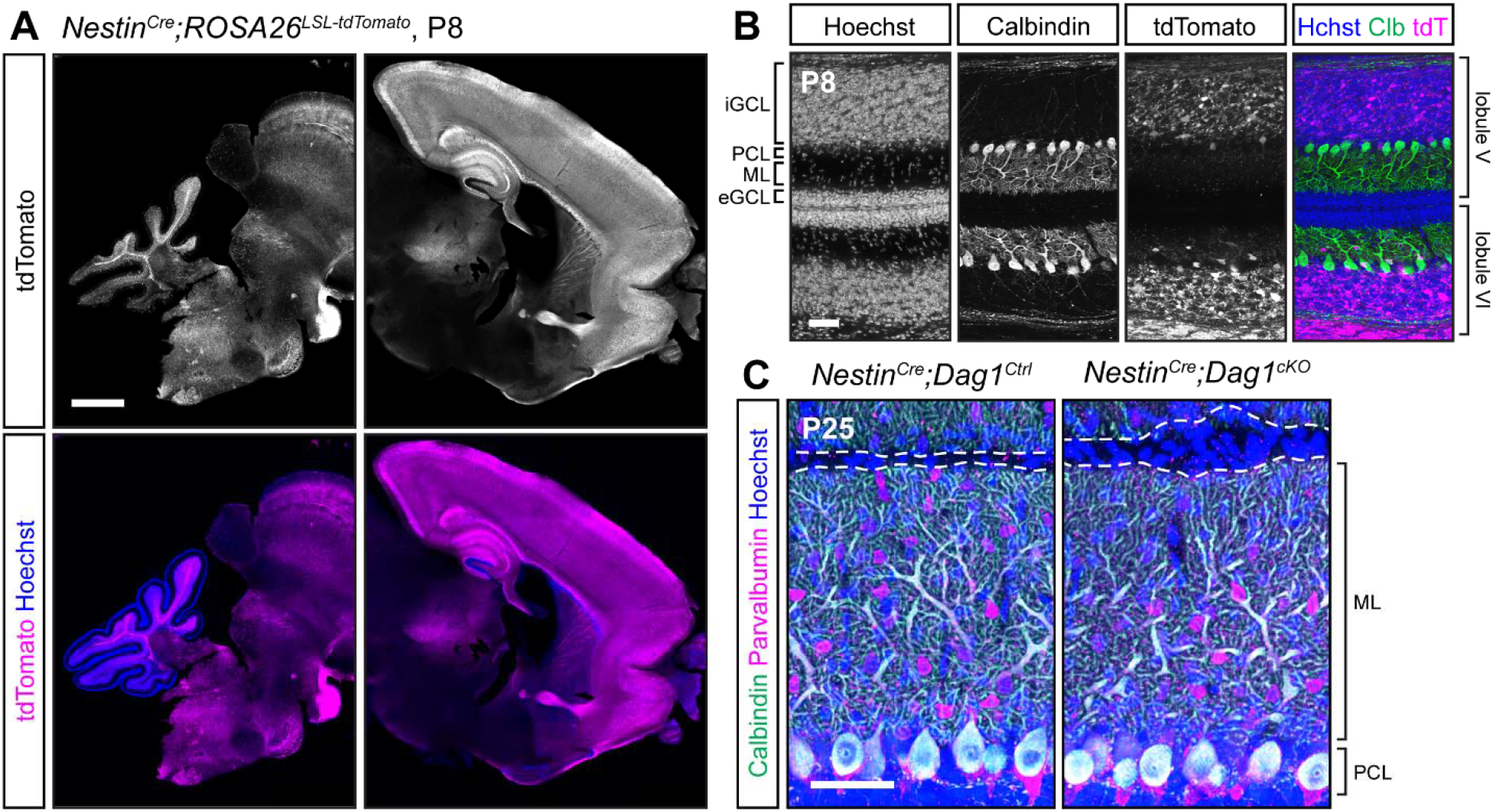
*Nestin^Cre^*does not drive *Cre* recombination in cerebellar Purkinje cells. **(A)** Fluorescent tdTomato reporter (magenta) in a sagittal brain slice of a P8 *Nestin^Cre^;ROSA^LSL-tdTomato^* mouse. Nuclei are stained in blue. Right and left panels are of the same image but adjusted differently as fluorescence in the forebrain is much brighter than the rest of the brain. Scale bar = 1000μm. **(B)** Higher magnification view of tdTomato expression pattern in cerebellar lobules V and VI at P8 (magenta). Purkinje cells are immunolabeled with anti-Calbindin (green). Scale bar = 50μm. (eGCL = external granule cell layer, iGCL = internal granule cell layer, PCL = Purkinje cell layer, ML = molecular layer.) **(C)** P25 *Nestin^Cre^;Dag1^cKO^* and littermate control Purkinje cells in cerebellar lobule V. Calbindin (green) labels Purkinje cells, Parvalbumin (magenta) labels Purkinje cells and molecular layer interneurons, Hoechst (blue) labels nuclei. Dashed line shows the edge of Purkinje cell dendrites at the pial surface. Scale bar = 50μm.

### Embryonic expression of *Ptf1a^Cre^* in GABAergic precursors results in stochastic deletion of *Dag1* from Purkinje cells despite ubiquitous reporter expression

We next tested a *Ptf1a^Cre^* line, which expresses *Cre* from the endogenous *Ptf1a* locus. *Ptf1a* is a transcription factor that is required for GABAergic neuronal fate in cerebellum (Hoshino et al., 2005). Examination of *Ptf1a^Cre^;ROSA26^LSL-tdTomato^*brains at P0 showed strong tdTomato fluorescence in the nascent cerebellum which co-localized with Calbindin, a marker for Purkinje cells (**Figure 5A, B**). Examination of adult (P21) brains showed tdTomato in all Purkinje cells and MLIs (**Figure 5A, C**). The early expression of tdTomato suggested that this line may be useful for developmental deletion of synaptogenic genes in Purkinje cells, although with reduced cellular specificity. However, when we examined Dag1 protein localization at P21, we found that *Ptf1a^Cre^;Dag1^cKO^* Purkinje cells showed mosaic loss of Dag1 protein, despite all Purkinje cells being tdTomato positive (**Figure 5C**). This result was surprising, as we anticipated that the early expression of *Cre* in Purkinje cells would delete the *Dag1* allele before it is expressed at appreciable levels. This difference between robust tdTomato expression and mosaic *Dag1* deletion may be due to the transient expression of *Ptf1a* in GABAergic precursors in conjunction with the known sensitivity of the particular *ROSA26^LSL-tdTomato^*floxed locus in the *Ai14* line to *Cre*-mediated recombination (Jin & Xiang, 2019; Luo et al., 2020). In this situation, the transient Cre activity can potentially drive recombination of the reporter allele without driving recombination of the *Dag1* floxed allele. These results suggest that the *Ptf1a^Cre^* line is of limited utility for studying deletion of proteins in GABAergic cerebellar neurons, as the presence tdTomato is not predictive of loss of the targeted allele.

**Figure 5.**
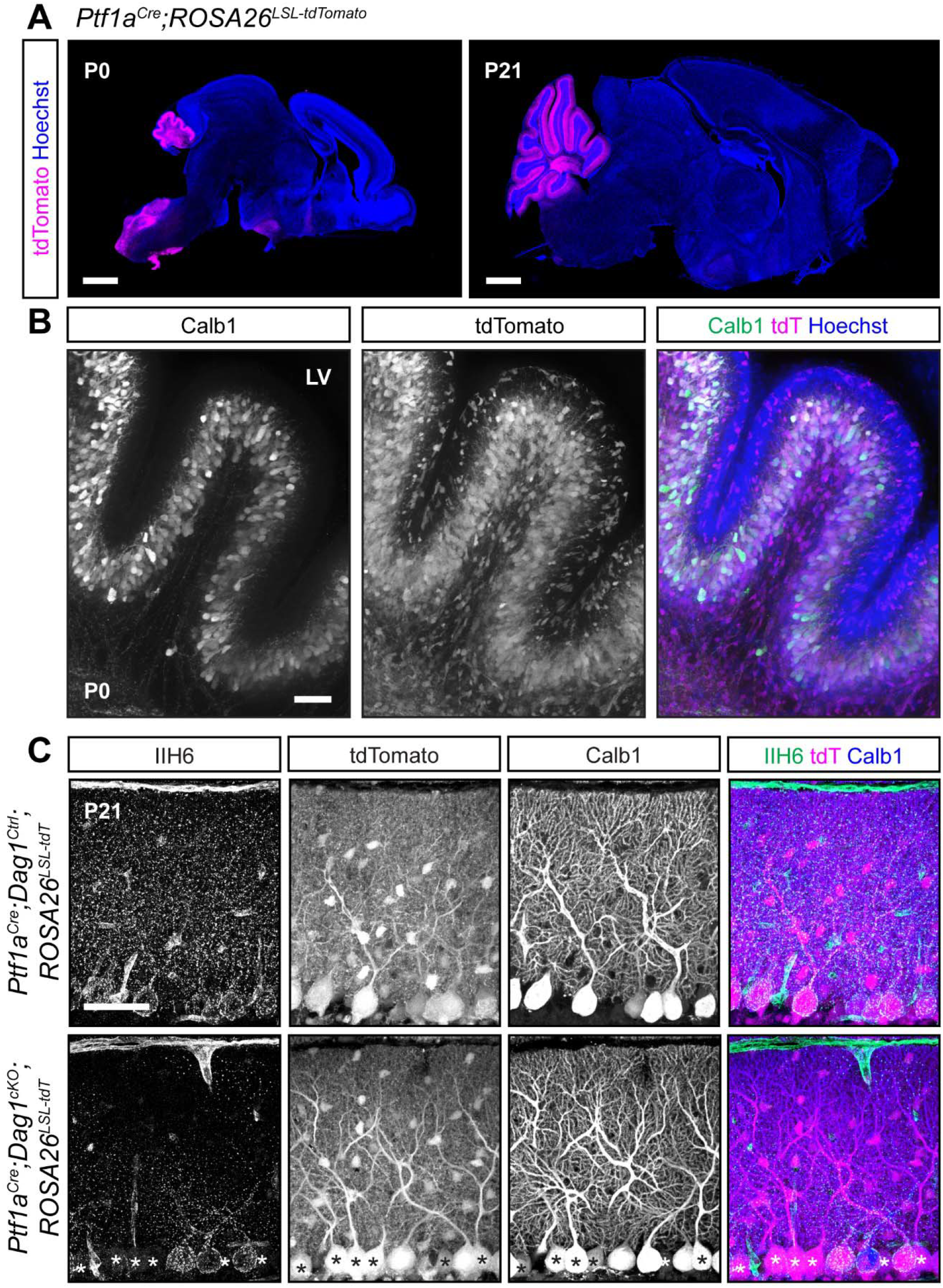
Early *Cre* recombination in developing Purkinje cells and MLIs with *Ptf1a^Cre^* results in a mosaic loss of Dystroglycan protein despite uniform reporter expression. (A) *Ptf1a^Cre^;ROSA26^LSL-tdTomato^*shows that *Cre* recombination, as reported by tdTomato (magenta), is evident at P0 and restricted to the cerebellum. Scale bars = 1000μm. **(B)** Higher magnification view of cerebellar lobule V at P0. Purkinje cells are labeled with Calbindin (Calb1, green) and *Cre* recombination is reported by native fluorescence of tdTomato (tdT, magenta). Nuclei are visualized with Hoechst (blue). Scale bar = 50μm. **(C)** Immunostaining in lobule V of P21 cerebella from *Ptf1a^Cre^;ROSA26^LSL-tdTomato^;Dag1^cKO^*and littermate controls for α-Dystroglycan (IIH6, green) to visualize Dag1 protein and Calbindin (Calb1, blue) to label Purkinje cells. *Cre* recombination was assessed by native fluorescence of the tdTomato reporter (tdT, magenta). Asterisks denote Purkinje cells with no detectable IIH6 signal. Scale bar = 50μm.

### *Calb1^Cre^* drives stable *Cre* expression and *Dag1* deletion in all Purkinje cells prior to synaptogenesis

In the course of characterizing the *Ptf1a^Cre^* line, we observed abundant Calbindin staining in Purkinje cells at P0 (**Figure 5**), and previous work has shown *Calb1* expression in the cerebellum as early as E14.5 (Morales & Hatten, 2006). While Calbindin is widely expressed in multiple neuronal subtypes in the majority of the CNS, within the cerebellum its expression is highly selective for Purkinje cells. *Calb1-IRES-Cre-D* mice (hereafter referred to as *Calb1^Cre^*) express *Cre* from the endogenous *Calb1* locus while retaining *Calb1* expression (Daigle et al., 2018). Analysis of *Calb1^Cre^;ROSA26^LSL-tdTomato^*brains showed tdTomato signal in all Purkinje cells at P0 (**Figure 6A**). Further analysis of staining at P6 and P14 continued to show that recombination is specific to Purkinje cells (**Figure 6B, C**). We next examined *Dag1* deletion in *Calb1^Cre^;Dag1^cKO^* mice at P14 during the period of active synaptogenesis in Purkinje cells. In contrast to the *Pcp2^Cre^;Dag1^cKO^*mice, which retained Dag1 at P16, we saw a complete loss of punctate Dag1 signal in Purkinje cells at P14 in *Calb1^Cre^;Dag1^cKO^* cerebella. These results highlight the utility of the *Calb1^Cre^* line to study the effect of gene knockout in Purkinje cells early in development.

**Figure 6.**
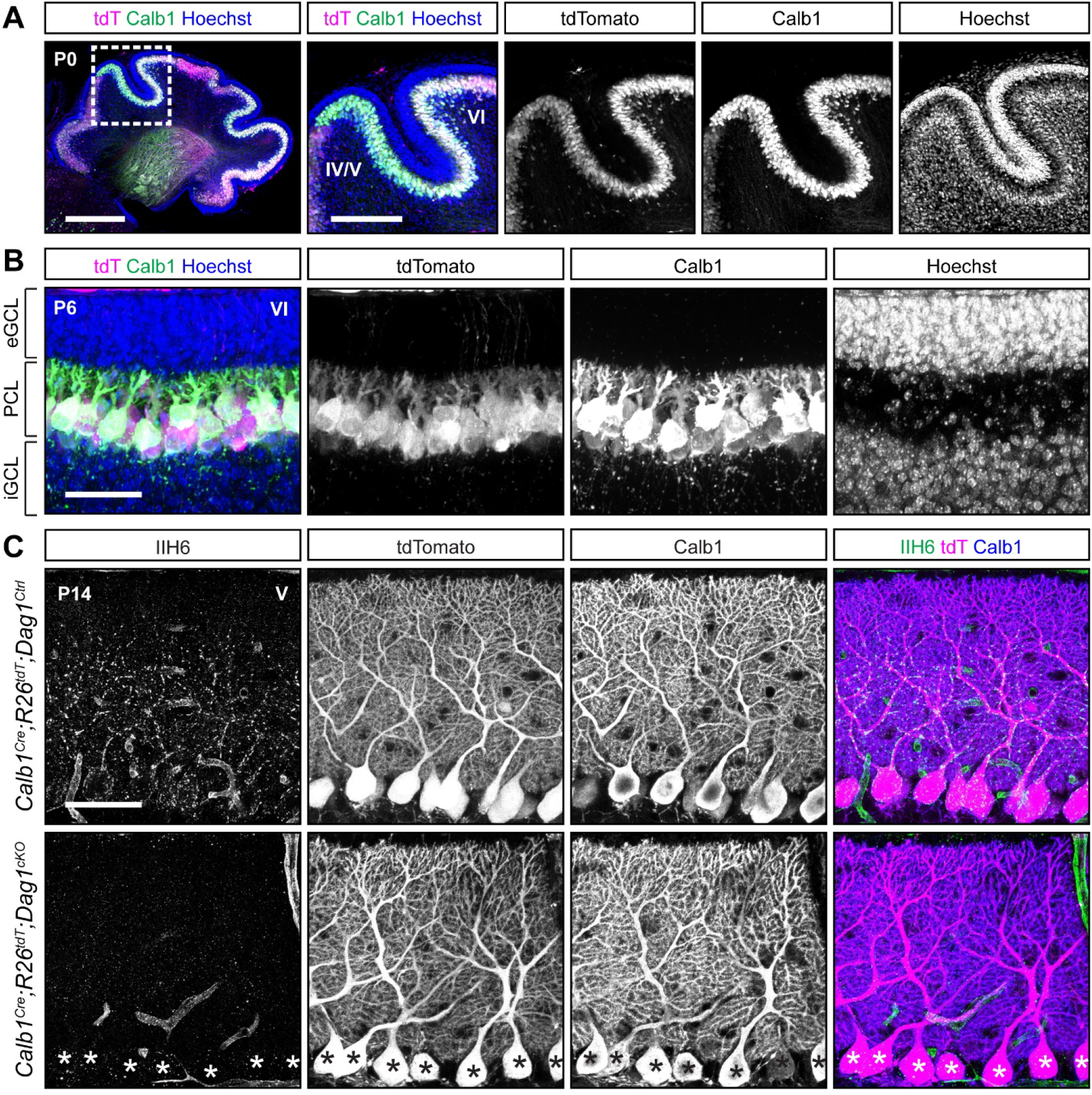
*Calb1^Cre^*drives *Cre* recombination early in development resulting in complete loss of synaptic Purkinje cell Dystroglycan protein. (A-B) Cerebellum of **(A)** P0 (insets show lobules IV/V and VI) and **(B)** P6 (lobule VI) *Calb1^Cre^;ROSA26^LSL-tdTomato^*mouse immunolabeled with Calbindin (Calb1, green) to label Purkinje cells and Hoechst (blue) to label nuclei. *Cre* recombination is reported by native tdTomato fluorescence (tdT, magenta). (eGCL = external granule cell layer, iGCL = internal granule cell layer, PCL = Purkinje cell layer.) Scale bars = **(A)** 500μm, 250μm (inset), **(B)** 50μm. **(C)** Immunostaining in lobule V of P14 cerebella from *Calb1^Cre^;ROSA26^LSL-tdTomato^;Dag1^cKO^*and littermate controls for α-Dystroglycan (IIH6, green) to visualize Dag1 protein and Calbindin (Calb1, blue) to label Purkinje cells. *Cre* recombination was assessed by native fluorescence of the tdTomato reporter (tdT, magenta). Asterisks denote Purkinje cells with no detectable IIH6 signal. Scale bar = 50μm.

### AAV delivery of *Cre* to Purkinje cells results in gradual *Cre* expression over the course of three weeks

In addition to transgenic *Cre* lines, viral-mediated delivery of *Cre* is commonly used to delete synaptogenic genes throughout the brain. One advantage of viral-mediated deletion is the ability to test cell-autonomous deletion of target alleles in a mosaic fashion by titrating the amount of virus. However, this approach is limited by the timing of viral transduction, subsequent *Cre* expression, and recombination. To better define this time course in Purkinje cells, we examined recombination of tdTomato in *ROSA26^LSL-tdTomato^* mice injected in the lateral cerebral ventricles at P0 with dilute *AAV8-CMV-Cre*. The first tdTomato positive Purkinje cells were observed around P7-P10, with the number of positive cells increasing to maximal levels by P18 (**Figure 7A, B**). This timing is similar to the recombination driven by *Pcp2^Cre^*, limiting its utility to examining synapse maintenance. Examination of Dag1 protein in *Dag^flox/-^;ROSA26^LSL-tdTomato^* mice injected with *AAV8-CMV-Cre* at P0 showed that loss of Dag1 lagged behind tdTomato expression, similar to what was observed in the *Pcp2^Cre^;Dag1^cKO^*Purkinje cells (**Figure 7C-F**). All *tdTomato*-expressing Purkinje cells were immunoreactive for Dag1 protein at P18, and most (90.8 ± 6.5%) remained Dag1 immunoreactive at P21 (**Figure 7C-D, F**). By P35 the majority of tdTomato expressing Purkinje cells lacked Dag1 immunoreactivity (**Figure 7E, F**). This lag between tdTomato expression and loss of Dag1 protein implies that it can take two or more weeks for Dag1 synaptic protein to turn over.

**Figure 7.**
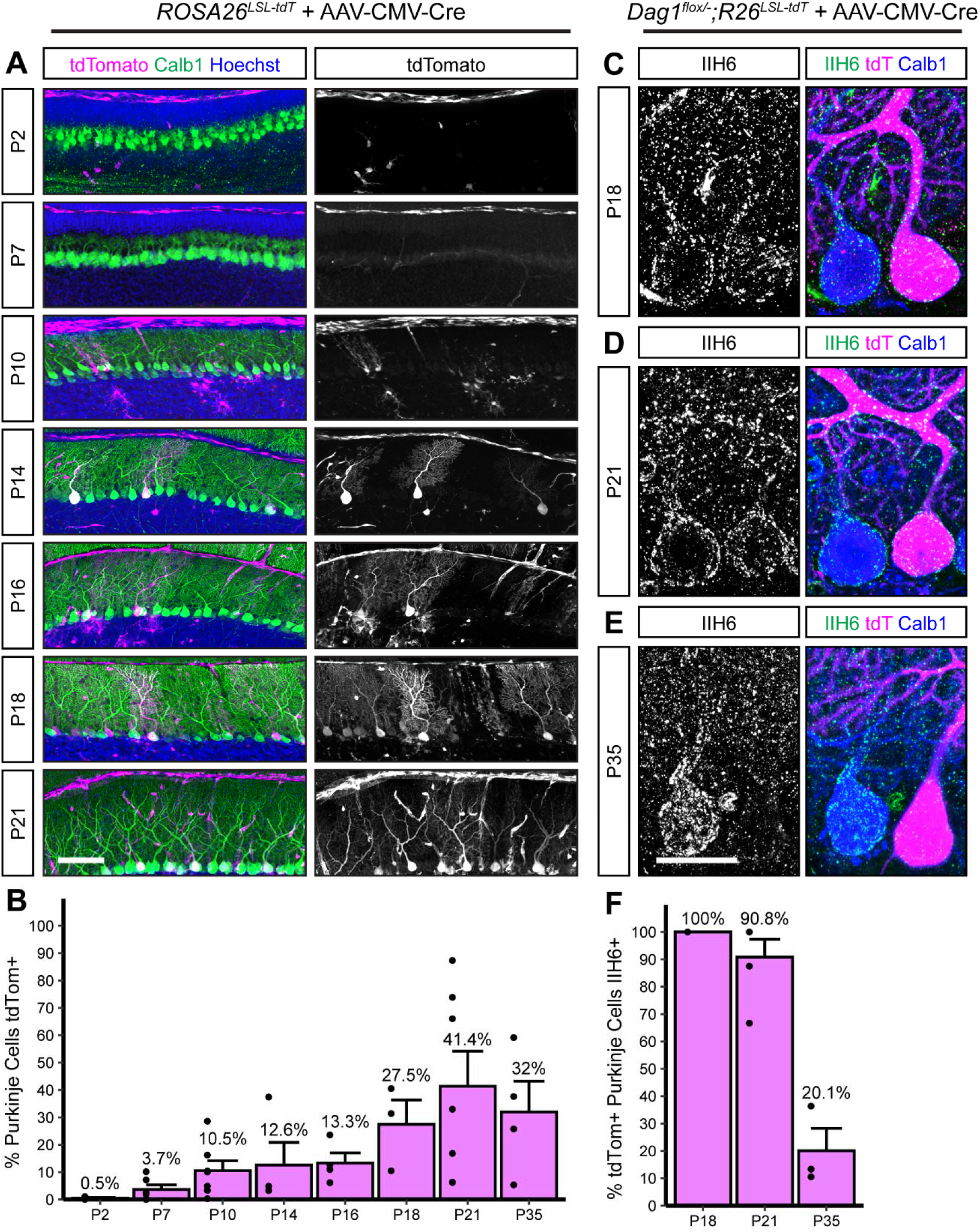
Viral delivery of *Cre* under the ubiquitous CMV promoter takes several weeks to recombine the *Dag1* floxed locus in Purkinje cells. (A) Dilute AAV8-CMV-Cre (1.00E+12) was injected into the lateral ventricles of P0 *ROSA26^LSL-tdTomato^*reporter mice. Cerebella were collected at timepoints from P2 to P35 to observe tdTomato expression. Sagittal sections were immunolabeled with Calbindin (Calb1, green) to label Purkinje cells and Hoechst (blue) to label nuclei. *Cre* recombination was assessed by native fluorescence of the tdTomato reporter (magenta). Scale bar = 100μm. **(B)** Quantification of the percent of Purkinje cells that express tdTomato in **(A)**. (C-E) *Dag1^flox/-^*;*ROSA26^LSL-tdTomato^*mice were injected with dilute AAV8-CMV-Cre at P0 and brains collected at **(C)** P18, **(D)** P21, and **(E)** P35. Cerebellar sections were counterstained with Calbindin (Calb1, blue) to label Purkinje cells and IIH6 (green) to label Dag1 protein. *Cre* expressing Purkinje cells were visualized with native tdTomato expression (magenta). Scale bar = 25μm. **(F)** Quantification of the percent of tdTomato expressing Purkinje cells that are IIH6 immunoreactive in **(C-E)**.

## Discussion

The Cre-lox system has proven to be an invaluable tool for studying the role of genes in a population-specific manner as conditional gene deletion is often required to avoid lethality of constitutive deletion. However, our results highlight the need for rigorous validation to ensure that a transgenic *Cre* line drives recombination and protein loss in expected cell types and time points. While the timing of *Cre* expression in a given line should be consistent across different conditional lines, the recombination efficiency of the floxed allele and the stability of existing protein that must be turned over before a cell can be deemed a “knockout” can vary. For these reasons, it is important to use a *Cre*-dependent reporter to evaluate specificity, along with a method for evaluating loss of protein (or mRNA) to validate deletion.

Each *Cre*-dependent reporter line may have different advantages and limitations. The *ROSA26* and *TIGRE* loci are popular sites for insertion of *Cre*-dependent fluorescent reporters due to their genomic accessibility. However, even reporters within the same loci (i.e. *ROSA26*) can show differences in recombination efficiency, which could lead to confusion about patterns of *Cre* expression (Daigle et al., 2018; Madisen et al., 2010). There have also been cases reported in which a *Cre* line may become lethal or cause health complications when crossed to certain fluorescent reporter lines but not others (Daigle et al., 2018). Here we used the popular *Ai14* line which expresses *LSL-tdTomato* in the *ROSA26* locus. The *Ai14* line is particularly sensitive to recombination, likely due to the proximity of the two loxP sites to one another. This could lead to off-target reporter expression and requires especially rigorous validation of any knockout when paired with inducible *Cre* lines which require administration of tamoxifen to activate the Cre protein (Álvarez-Aznar et al., 2020; Luo et al., 2020).

Inducible *Cre* lines (e.g. *CreER^T2^)* are often used to either control the timing of Cre activity or to restrict Cre activity to a subset of a cellular population. However, because of the transient nature, inducible *Cre* lines require especially careful validation. *CreER^T2^* consists of *Cre* fused to the hormone binding domain of the estrogen receptor (R. Feil et al., 1997; S. Feil et al., 2009). Under baseline conditions, CreER^T2^ localization is excluded from the nucleus. Once tamoxifen is administered it is metabolized into 4-hydroxytamoxifen, a synthetic ligand of the estrogen receptor that binds to CreER^T2^ and permits localization to the nucleus. After tamoxifen has been cleared from the system, CreER^T2^ is once again excluded from the nucleus, restricting CreER^T2^ activity to roughly 24 hours after tamoxifen administration (R. Feil et al., 1997). Since the effect of tamoxifen is both transient and dose-dependent, it is prudent that a *Cre*-dependent reporter is used to identify which cells experienced Cre-mediated recombination. However, due to differences in sensitivity to recombination between floxed alleles, reporter expression cannot be used alone to predict loss of protein (S. Feil et al., 2009).

Based on expression of tdTomato driven by *Pcp2^Cre^*, we observed that *Pcp2^Cre^* drives *Cre* expression selectively but gradually in Purkinje cells beginning at P7 and reaching all Purkinje cells by P14 (**Figure 2**). This time course is similar to what has been reported for the related *L7^Cre^* line (Barski et al., 2000; Lewis et al., 2004). However, when *L7^Cre^* was used to delete *Dag1* from Purkinje cells, loss of protein wasn’t complete until P90 (Briatore et al., 2020), whereas we observed loss of Dag1 protein by P30 (**Figure 3**). This difference in the loss of protein is likely due to differing breeding strategies. Briatore and colleagues report crossing *L7^Cre^;Dag1^flox/flox^*mice with *Dag1^flox/flox^* mice to generate *L7^Cre^;Dag1^flox/flox^*mutants and *Dag1^flox/flox^* littermate controls. While this is an efficient breeding strategy allowing for the use of all progeny in a litter, the use of flox/flox mutants may not be ideal for studying *Dag1* specifically. Dag1 is a stable protein, and in this case *Cre* must recombine both alleles of *Dag1* before protein loss can begin. We crossed *Pcp2^Cre^;Dag1^+/-^* mice to *Dag1^flox/flox^* mice to generate *Pcp2^Cre^;Dag1^flox/-^*mutants and *Pcp2^Cre^;Dag1^flox/+^* littermate controls. Not only does this approach control for potential off-target effects of *Cre* expression by ensuring that both controls and mutants are *Pcp2^Cre^*expressing, *Cre* only needs to recombine one floxed allele of *Dag1*. This alone could account for the faster loss of protein. However, this approach requires ensuring that heterozygous control mice lack a phenotype. While *L7^Cre^*and *Pcp2^Cre^* remain useful tools for studying the effect of losing protein later in development, an inducible *Cre* such as *Pcp2^CreERT2^*with tamoxifen administration at or after P14 might be more useful for standardizing the timing of gene deletion across Purkinje cells, avoiding gradual *Cre* expression seen in the *Pcp2^Cre^* line. However, as described above, tamoxifen administration represents a new variable with additional optimization and validation required.

The time course of *Cre* expression in Purkinje cells with intracerebroventricular delivery of *AAV8-CMV-Cre* at P0 was similar to that observed with *Pcp2^Cre^*(**Figures 1, 7**). One advantage of viral delivery is the ability to achieve sparse or mosaic *Cre* expression. This can also be achieved using inducible *CreER^T2^*lines with low-dose tamoxifen; however virally-expressed *Cre* remains expressed in the cell after transduction whereas Cre activity with *CreER^T2^*is temporally limited to approximately 24 hours after tamoxifen administration (R. Feil et al., 1997). This increased stability from virally-expressed *Cre* is therefore less likely to result in expression of a *Cre*-dependent reporter without recombination of target floxed alleles, which can occur following transient CreER^T2^ activity induced by low doses of tamoxifen. The timing of AAV8 viral transduction limits the utility of this approach to later developmental processes. The use of AAVs for gene transduction has become standard in the field, and new viral capsids to expand the utility of the approach are under rapid development. Different capsids can exhibit differences in tropism, efficiency, and transduction timeline (Haery et al., 2019). While the most widely used capsids show a delay in transduction of several days to a couple weeks, newer developments can speed up that timeline considerably. A recent study identified an *AAV-SCH9* serotype that can transduce neurons in 24-48 hours, making it useful for studying developmental processes (Zheng et al., 2023).

Development of genetic tools for targeting inhibitory populations of neurons has lagged behind those available for targeting excitatory populations. Within the cerebellum Purkinje cells (the primary output of the cerebellum) and MLIs (local interneurons) are both GABAergic, making it challenging to separate the two based on common markers for inhibitory populations (ex. *VGAT^Cre^, GAD2^Cre^*). Both populations also express Parvalbumin, meaning *PV^Cre^* will cause recombination in both PCs and MLIs. *Dlx5/6^Cre^* is becoming increasingly popular to target forebrain interneuron populations, however *Dlx5/*6 is not expressed in the cerebellum. Here we have described a suite of tools for targeting Purkinje cells for either genetic labeling or gene deletion at various developmental stages: *Calb1^Cre^*for embryonic deletion (**Figure 6**), *Pcp2^Cre^* for early postnatal deletion (**Figures 2, 3**), and *AAV-Cre* for deletion during or after postnatal development (**Figure 7**). We also highlight important considerations when identifying and validating new tools for conditional genetic deletion applicable to any population of cells.

## Acknowledgements

This work was funded by NIH Grants R01NS091027 and R01NS131299 (KMW), F31NS120649 (JNJ), P30NS061800 (OHSU ALM) and CureCMD (KMW).

